# Escher-FBA: A web application for interactive flux balance analysis

**DOI:** 10.1101/281600

**Authors:** Elliot Rowe, Bernhard O. Palsson, Zachary A. King

## Abstract

**Background:** Flux balance analysis (FBA) is a widely-used method for analyzing metabolic networks. However, most existing tools that implement FBA require downloading software and writing code. Furthermore, FBA generates predictions for metabolic networks with thousands of components, so meaningful changes in FBA solutions can be difficult to identify. These challenges make it difficult for beginners to learn how FBA works.

**Results:** To meet this need, we present Escher-FBA, a web application for interactive FBA simulations within a pathway visualization. Escher-FBA allows users to set flux bounds, knock out reactions, change objective functions, upload metabolic models, and generate high-quality figures without downloading software or writing code. We provide detailed instructions on how to use Escher-FBA to replicate several FBA simulations that generate real scientific hypotheses.

**Conclusions:** We designed Escher-FBA to be as intuitive as possible so that users can quickly and easily understand the core concepts of FBA. The web application can be accessed at https://sbrg.github.io/escher-fba.

## Background

The constraint-based modeling approach to studying metabolic networks has led to a great variety of applications, from understanding metabolic gene essentiality, stress tolerance, and gene regulation to designing microbial cell factory [1]. The simplest and most popular constraint-based method is flux balance analysis (FBA) [2], and many derivative approaches draw their value from extending the elegant insights of FBA [3]. Most tools for FBA simulation require software downloads, have significant learning curves, and require computer programming. However, FBA has broad interest, so there is a great value in simple tools for FBA that can be picked up quickly by new users.

The metabolic networks that are used for FBA simulations are represented by genome-scale models (GEMs). GEMs are available for many model organisms, with the number growing rapidly [4]. Standard formats with clear specifications and workflows for creating high quality GEMs are available [5]. Any new tool for FBA needs to support import of the many existing models. Genome-scale models of key model organisms have many thousands of metabolites and reactions [6–8], so it can be very difficult for newcomers and experienced modelers to recognize the changes in FBA simulations. This is where visualizations of GEMs and associated data play a critical role in the modeling process [1].

The most popular software packages for FBA simulation require computer programming skills. COBRA Toolbox [9] and COBRApy [10] require knowledge of the MATLAB and Python programming languages, respectively. PSAMM is a another Python-based toolbox [11]. The only popular application for FBA that does not require computer programming is OptFlux [12]. Therefore, OptFlux has become popular as a teaching tool for FBA and other strain design algorithms. OptFlux and COBRA Toolbox both have FBA visualization features, and there are dedicated visualization tools like FBASimVis [13] and FluxViz [14] available. However, the best way to understand FBA is to explore simulations interactively by modifying parameters and receiving immediate feedback. Until now, no software has enabled this kind of interactive exploration of FBA simulations, and no existing FBA tools are fully web-based.

Escher-FBA meets these needs by extending the Escher application for pathway visualization with on-the-fly FBA calculations. Escher-FBA adds functionality so users can modify parameters in the FBA simulation—including flux bounds, objective function, and reaction knockouts—with immediate visualization of the results. Escher-FBA constructs the network and reaction data using the same input files as Escher. Additionally, since Escher-FBA is a web application, it works across platforms, including mobile devices. These features make Escher-FBA an ideal choice for academic labs and classrooms where students and researchers need a cross-platform visualization tool for learning and exploring FBA simulations.

## Implementation

Escher-FBA is built upon Escher [15], a versatile and user-friendly visualization tool for metabolic pathways. Users can quickly and easily create their own maps by first loading a GEM containing all the reactions in the system and then building visualizations comprised of both reactions (symbolized by arrows) and metabolites (symbolized by circles). Users can also load, modify, and save their maps, as well as maps that have been created by the community. Escher maps are stored as JavaScript Object Notation (JSON) files.

Escher-FBA extends Escher with interactive tooltips that appear when a user hovers the mouse over (or taps on a touchscreen) any reaction in the pathway visualization (Fig. 1). These tooltips contain controls that immediately modify the parameters of the FBA simulation. FBA simulations are executed using the GNU Linear Programming Kit (GLPK), which has been compiled to JavaScript and runs in the browser (https://github.com/hgourvest/glpk.js). The slider control in the tooltip adjusts both the upper and lower flux bounds of the reaction, while also displaying the current flux through the reaction. Value fields for upper and lower flux bounds allow the users to enter precise values. A **Knockout** button can be used to simulate a knockout of the associated reaction by setting both the upper and lower bounds of the reaction to zero. There is also a **Reset** button to reset the bounds to their original values from the loaded model. Finally, the **Maximize** and **Minimize** buttons can be used to change the FBA objective function to maximize or minimize the flux through the current reaction. The current objective and flux through that objective are displayed along the bottom-left corner of the screen, and buttons to reset the entire map back to its original data and to view a help menu are in the bottom-right corner of the screen. Perturbing the system with these controls produces immediately visible effects within the system. There is no need to manually re-run the simulation as Escher-FBA produces a new solution in response to user input.

**Figure 1:**
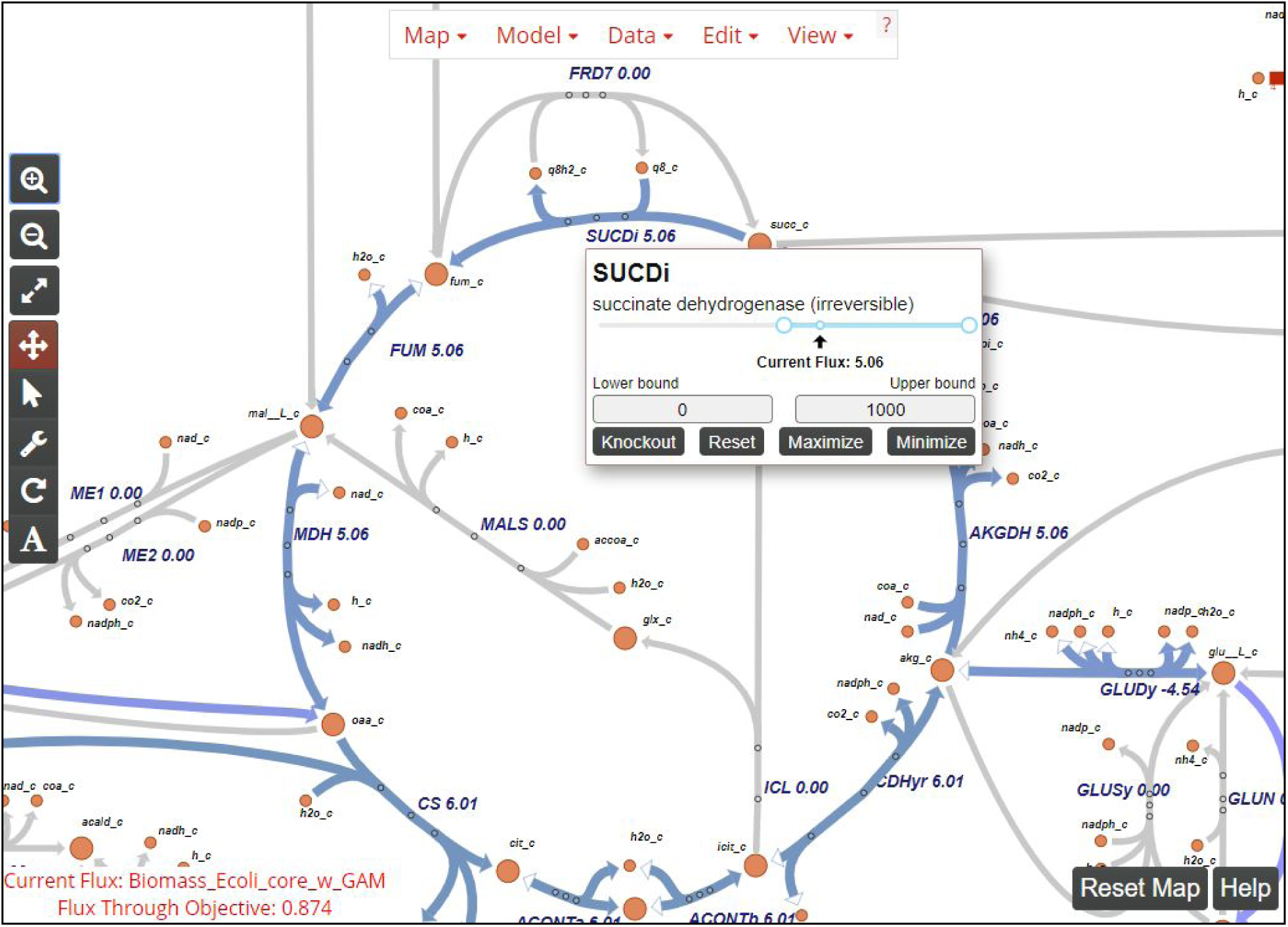
A screenshot of Escher-FBA. Buttons inside the tooltip for each reaction allow for quick modifications to be made to the network by the user with immediate visual updates to the network. At the bottom of the screen the current objective function, flux through the objective, and a reset button for the whole map can be seen.

Escher-FBA supports upload of custom maps and models, using the same upload functionality and file format as Escher [15]. Users can create their own maps and models and perform *in silico* experiments with their data. Additional maps and models can be downloaded from BiGG Models (http://bigg.ucsd.edu) [16]. The default map and model in Escher-FBA is a core model of central glucose metabolism in *E. coli* K-12 MG1655 (available at http://bigg.ucsd.edu/models/e_coli_core). This model is small enough that the user can see everything happening in the simulation within a single pathway map. Escher-FBA also supports full GEMs. Any GEM in the COBRA JSON file format can be imported in Escher-FBA directly. Models in other formats can be converted to JSON using COBRApy [10]. COBRApy supports many other file formats, including the latest Systems Biology Markup Language (SBML) with the Flux Balance Constraints (FBC) extension [17]. The JSON specification for COBRA models is also being adopted by other tools [11].

## Results and discussion

In order to demonstrate the use of Escher-FBA for real applications, we present four key FBA examples that can be executed directly in the browser. These are adapted from a review of FBA and its applications [2]. These examples rely on the default core model of *E. coli*, so they are ready to be implemented as soon as the Escher-FBA webpage is opened. Make sure to click the **Reset Map** button between each example. If you are having trouble finding a reaction, simply click the **Find** option in the **View** menu (or the “f” key on your keyboard) to open up a search bar.

The first example demonstrates the use of FBA to predict whether growth can occur on alternate carbon substrates. The default core model of *E. coli* includes a simulated minimal medium with D-glucose as the carbon source. Here, we will switch the carbon source from D-glucose to succinate. First, mouse over the succinate exchange reaction **EX_succ_e**, and change the lower bound to −10 mmol/gDW/hr, either by dragging the slider or by entering −10 into the **Lower Bound** field. Next, mouse over the D-glucose exchange reaction **EX_glc_e**, and either raise the lower bound to 0 or click the **Knockout** button. The default objective is still to maximize growth, so these two changes will instruct the program to calculate the maximum growth rate while using succinate as the carbon source instead of D-glucose. You should see that the maximum predicted growth rate decreases from 0.874 hr^−1^ to 0.398 hr^−1^, reflecting the lower growth yield of *E. coli* on succinate (Fig. 2a). This is the general approach to making changes in Escher-FBA; mouse over the reaction, make the required changes, and Escher-FBA will automatically display your results. The lower bound values for carbon source exchange represent experimental measurements, so you can try adjusting the specific lower bound value to realistic values for growth on other carbon sources.

**Figure 2:**
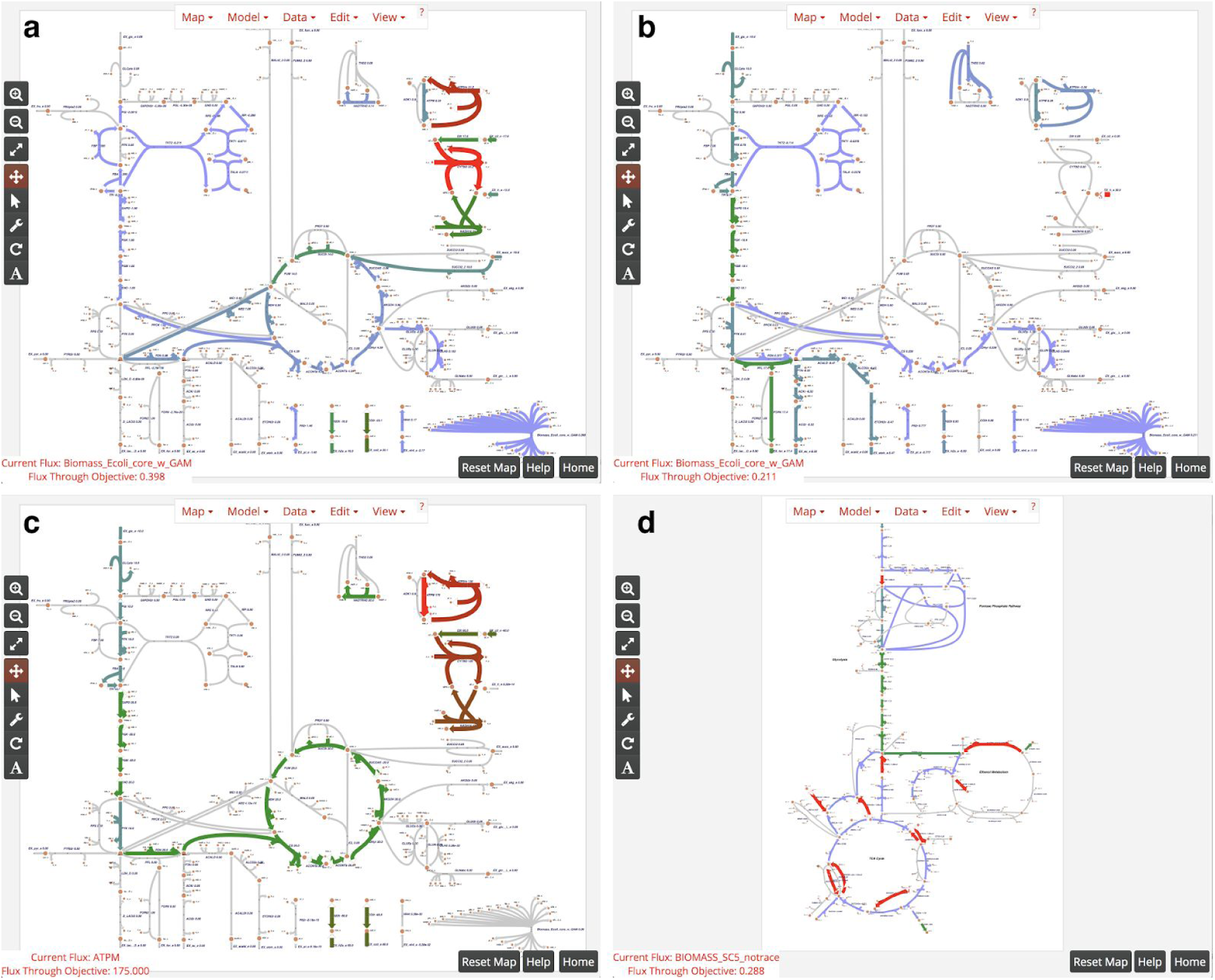
Examples of Escher-FBA simulations. (a) Simulated growth with succinate as sole carbon source. (b) Simulated anaerobic growth on a glucose minimal medium. (c) Maximizing ATP yield in the default model. (d) Growth of the *i*MM904 model of *S. cerevisiae*. Note that arrow widths were increased in the settings menu to make changes more obvious.

Anaerobic growth can be simulated in the same way by mousing over the **EX_o2_e** reaction and either clicking **Knockout** or change the lower bound to 0. If you change oxygen exchange to zero while succinate is still the only only carbon source, the **Flux Through Objective** indicator displays “Infeasible solution/Dead cell”, meaning that growth is not possible. Try clicking the **Reset** button in the bottom right corner to simulate a minimal medium with D-glucose as a carbon source, then knock out **EX_o2_e**, and the predicted growth rate should be 0.211 hr^−1^ (Fig. 2b).

We can also use Escher-FBA to determine the maximum yields of precursors and cofactors such as ATP. All that is required is a stoichiometrically balanced reaction that consumes the cofactor of interest. The ATP Maintenance (**ATPM**) reaction is one such example. In order to determine the maximum production of ATP, simply mouse over the **ATPM** reaction and click the **Maximize** button. Setting up the objective this way works because, in order for the system to maximize flux through the **ATPM** reaction, it must first produce ATP in the highest possible quantity. When **ATPM** is maximized in the default core metabolism model of *E. coli*, the objective value is 175 mmol/gDW/hr (Fig. 2c). With succinate as a carbon source, this value decreases to 82.5 mmol/gDW/hr. This same procedure can be followed for any metabolite of interest by creating a stoichiometrically-balanced consumption reaction and setting the model to maximize the flux through that reaction. Note that it is not currently possible to create such a reaction automatically in Escher-FBA, but this can be added to a future release.

Analyzing alternate optimal solutions in metabolism is another useful application of FBA [18]. Since the solutions produced through FBA are often non-unique, it can be useful to know the range of flux values a particular reaction can have. Flux variability analysis (FVA) is often used to calculate these ranges across the entire network [19]. Escher-FBA does not support FVA calculations directly, but it is possible to calculate them for a given reaction. In order to do this, first mouse over the objective function (the biomass reaction **Biomass_Ecoli_core_w_GAM**) and set the upper and lower bounds to slightly less than the current flux value (in the default map, try 0.870). Next, mouse over a reaction of interest and click the **Maximize** and **Minimize** buttons to see the maximum and minimum flux through that reaction given the optimal growth rate. For example, maximizing and minimizing flux through **GAPD** in glycolysis yields a feasible flux range of 15.44–16.68 mmol/gDW/hr, indicating that glycolytic flux is highly constrained at high growth rates. On the other hand, maximizing and minimizing flux through **MALS** in the glyoxylate shunt yields a feasible flux range of 0–2.64 mmol/gDW/hr, indicating that the glyoxylate shunt can be activate or inactive at high growth rates. This procedure can be done with any set of reactions and the user can constrain their system to any number of flux values to see the range of solutions available to a particular reaction.

The default *E. coli* core model is not the only system that can be simulated. For example, if one wishes to run simulations on a yeast cell, a model and map for *Saccharomyces cerevisiae* can be downloaded from http://bigg.ucsd.edu/models/iMM904. On that page, click the download button for the model (iMM904.json) and the map (iMM904.Central carbon metabolism.json). Load these in Escher-FBA by clicking **Load Map JSON** in the **Map** menu and **Load Model JSON** in the **Model** menu to load both JSON files. Once loaded, the map is ready to edit and simulate with any of the tools in Escher or Escher-FBA (Fig. 2d). With a larger model like iMM904, not all reactions will be visible at once, but you can add a reaction to the visualization. First either click the wrench icon on the sidebar or select **Add reaction mode** from the **Edit** menu. Now, reactions can be added by clicking anywhere on the map and selecting the desired reaction from the drop down menu. The text input field can be used to search for a reaction of interest.

While Escher-FBA can already be used for many FBA simulations directly in the web browser, a number of the examples presented by Orth *et al.* cannot currently be accomplished with Escher-FBA [2]. As of right now, Escher-FBA cannot perform functions such as gene knockout analysis or robustness analysis. However, Escher-FBA uses flexible SVG representations for visual elements, so FVA, robustness analysis, and even graphical features such as phase planes could be added.

## Conclusions

Escher-FBA can perform interactive FBA simulations without any software installation or knowledge of computer programming. Additionally, even though it currently cannot perform the advanced analysis available in popular libraries like the COBRA Toolbox and COBRApy, Escher-FBA can serve as an entrypoint for new users of FBA, for teaching FBA to students, and as a quick reference to experts who want to visually explore simulations.

## Availability and requirements

**Project name:** Escher-FBA

**Project home page:** https://sbrg.github.io/escher-fba

**Operating system:** Platform independent

**Programming language:** JavaScript

**Any restrictions to use by non-academics:** none

## List of abbreviations

FBA: Flux Balance Analysis
FVA: Flux Variability Analysis
GEM: Genome-scale Model
JSON: JavaScript Object Notation
SBML: Systems Biology Markup Language FBC - Flux Balance Constraints
GLPK: GNU Linear Programming Kit

## Declarations

### Ethics approval and consent to participate

Not applicable

### Consent for publication

Not applicable

### Availability of data and material

The application and documentation are available at https://sbrg.github.io/escher-fba.

### Authors’ contributions

ER and ZAK designed and implemented the application. ER, BOP, and ZAK wrote the manuscript.

## Acknowledgements

Special thanks to Niko Sonnenschein and Siddharth Chauhan for their feedback during the development of Escher-FBA.

